# Structural impact on SARS-CoV-2 spike protein by D614G substitution

**DOI:** 10.1101/2020.10.13.337980

**Authors:** Jun Zhang, Yongfei Cai, Tianshu Xiao, Jianming Lu, Hanqin Peng, Sarah M. Sterling, Richard M. Walsh, Sophia Rits-Volloch, Piotr Sliz, Bing Chen

## Abstract

Substitution for aspartic acid by glycine at position 614 in the spike (S) protein of severe acute respiratory syndrome coronavirus 2 (SARS-CoV-2), the causative agent of the ongoing pandemic, appears to facilitate rapid viral spread. The G614 variant has now replaced the D614-carrying virus as the dominant circulating strain. We report here cryo-EM structures of a full-length S trimer carrying G614, which adopts three distinct prefusion conformations differing primarily by the position of one receptor-binding domain (RBD). A loop disordered in the D614 S trimer wedges between domains within a protomer in the G614 spike. This added interaction appears to prevent premature dissociation of the G614 trimer, effectively increasing the number of functional spikes and enhancing infectivity. The loop transition may also modulate structural rearrangements of S protein required for membrane fusion. These findings extend our understanding of viral entry and suggest an improved immunogen for vaccine development.

## Introduction

Severe acute respiratory syndrome coronavirus 2 (SARS-CoV-2), an enveloped positive-stranded RNA virus, is the cause of the ongoing COVID-19 pandemic. It probably originated when related viruses circulating in bats acquired the ability to infect human cells^1^. While viral evolution is believed to be slow, owing to the RNA proofreading capability of its replication machinery^2^, a variant with a single-residue substitution (D614G) in its spike protein has rapidly become the dominant strain throughout the world^3^. Understanding the molecular features of the now most prevalent virus strain can guide intervention strategies to control the crisis.

SARS-CoV-2 initiates infection by fusion of its envelope lipid bilayer with the membrane of a host cell. This critical step in the viral life cycle is catalyzed by the trimeric spike (S) protein, which is produced as a single-chain precursor and subsequently processed by a furin-like protease into the receptor-binding fragment S1 and the fusion fragment S2^4^. After engagement of the receptor-binding domain (RBD) in S1 with the viral receptor angiotensin converting enzyme 2 (ACE2) on the host cell surface, followed by a second proteolytic cleavage within S2 (S2’ site)^5^, the S protein undergoes large conformational changes, prompting dissociation of S1 and irreversible refolding of S2 into a postfusion structure^6,7^. Formation of the postfusion S2 provides energy for overcoming the kinetic barrier of membrane fusion and effectively brings the viral and cellular membranes close together to induce fusion of the two.

Rapid advances in the structural biology of the SARS-CoV-2 S protein, an important target for development of diagnostics, therapeutics and vaccines, include structures of S protein fragments derived from the original virus carrying D614: the S ectodomain stabilized in its prefusion conformation^8,9^, RBD-ACE2 complexes^10-13^, and segments of S2 in the postfusion state^14^. In the prefusion ectodomain structure, S1 folds into four domains - NTD (N-terminal domain), RBD, and two CTDs (C-terminal domains), and wraps around the prefusion conformation of S2, with the RBD sampling two distinct conformations – “up” for a receptor-accessible state and “down” for a receptor-inaccessible state. We and others have also reported structures of a purified, full-length D614 S protein in both prefusion and postfusion conformations^15,16^. Studies by cryoelectron tomography, with chemically inactivated SARS-CoV-2 preparations, using both D614 and G614 variants have revealed additional structural details of S proteins present on the surface of virion^17-20^.

Epidemiological surveillance indicated that the SARS-CoV-2 carrying G614 outcompeted the original virus and became the globally dominant form within a month^3,21,22^. This single-residue substitution appears to correlate with high viral loads in infected patients and high infectivity of pseudotyped viruses, but not with disease severity^3^. The G614 virus has comparable (or even slightly higher) sensitivity to neutralization by convalescent human sera or vaccinated hamster sera^3,23-25^, suggesting that the response to vaccination with immunogens containing D614 remains effective against the new strain. Further studies have demonstrated that S1 dissociates more readily from the D614 virus than from G614 virus^26^, indicating that the D614 viral spike is substantially less stable than the G614 variant. The stabilized, soluble S ectodomain trimer with G614 samples the RBD-up conformations more frequently than does the D614 trimer^25,27^, but it is puzzling why the former binds more weakly to recombinant ACE2 than the latter^27^. The known S trimer structures indicate that the D614G change breaks a salt bridge between D614 and a positively charged residue (K854) in the fusion peptide proximal region (FPPR)^15^, which may help clamp the RBD in the prefusion conformation. This observation can explain why the G614 trimer favors the RBD-up conformations, but does not account for its increased stability. To resolve these issues, which relate directly to viral entry mechanism and to vaccine development, we report here the structural consequences of the D614G substitution in the context of the full-length S protein.

## Results

### Characteristics of the full-length SARS-CoV-2 S protein carrying G614

Following our established protocols for producing the full-length S protein containing D614 solubilized in detergent^15^, we transfected HEK293 cells with a construct expressing a full-length wildtype SARS-CoV-2 S with G614. We compared the membrane fusion activity of the G614 S protein with that of the full-length D614 S construct in a β-galactosidase-based cell-cell fusion assay^15^. As shown in Fig. S1A, all the cells expressing S fused efficiently with cells transfected with a human ACE2 construct, demonstrating that the S proteins expressed on the cell surfaces are fully functional. At the low transfection levels, the G614 S had higher fusion activity than the D614 S, but the difference diminished with the increased amount of transfected DNA, suggesting that the high expression levels can compensate for any possible defects that associated with the D614 S protein. We also tested inhibition of the cell-cell fusion by an engineered trimeric ACE2-based inhibitor that competes with the receptor on the target cells^28^, showing that the G614 trimer is slightly more sensitive than the D614 trimer (Fig. S1B).

To purify the full-length S protein, we used an expression construct fused with a C-terminal strep-tag, which was equally active in cell-cell fusion as the untagged version (Fig. S1A), and purified both G614 and D614 proteins under identical conditions. We lysed the transfected cells and solubilized membrane-bound proteins in 1% detergent dodecyl-β-D-maltopyranoside (DDM), purified the strep-tagged S proteins by elution from strep-tactin resin in 0.3% DDM followed by gel filtration chromatography in 0.02% DDM. The D614 protein eluted in three peaks representing the prefusion S trimer, the postfusion S2 trimer and the dissociated monomeric S1, respectively, as we have reported previously^15^. The G614 protein eluted as a single major peak, corresponding to the prefusion S trimer (Fig. 1A). Coomassie-stained SDS-PAGE analysis confirmed that the G614 peak contained mainly the cleaved S1/S2 complex (~90%) and a small amount of the uncleaved S precursor (~10%). We found similar patterns when the two proteins were purified in detergent NP-40, indicating that the choice of detergent had not affected the relative stability of the two spike variants. We conclude that the single-residue substitution has a striking effect on the stability of the SARS-CoV-2 S trimer, even as a purified protein.

**Figure 1.**
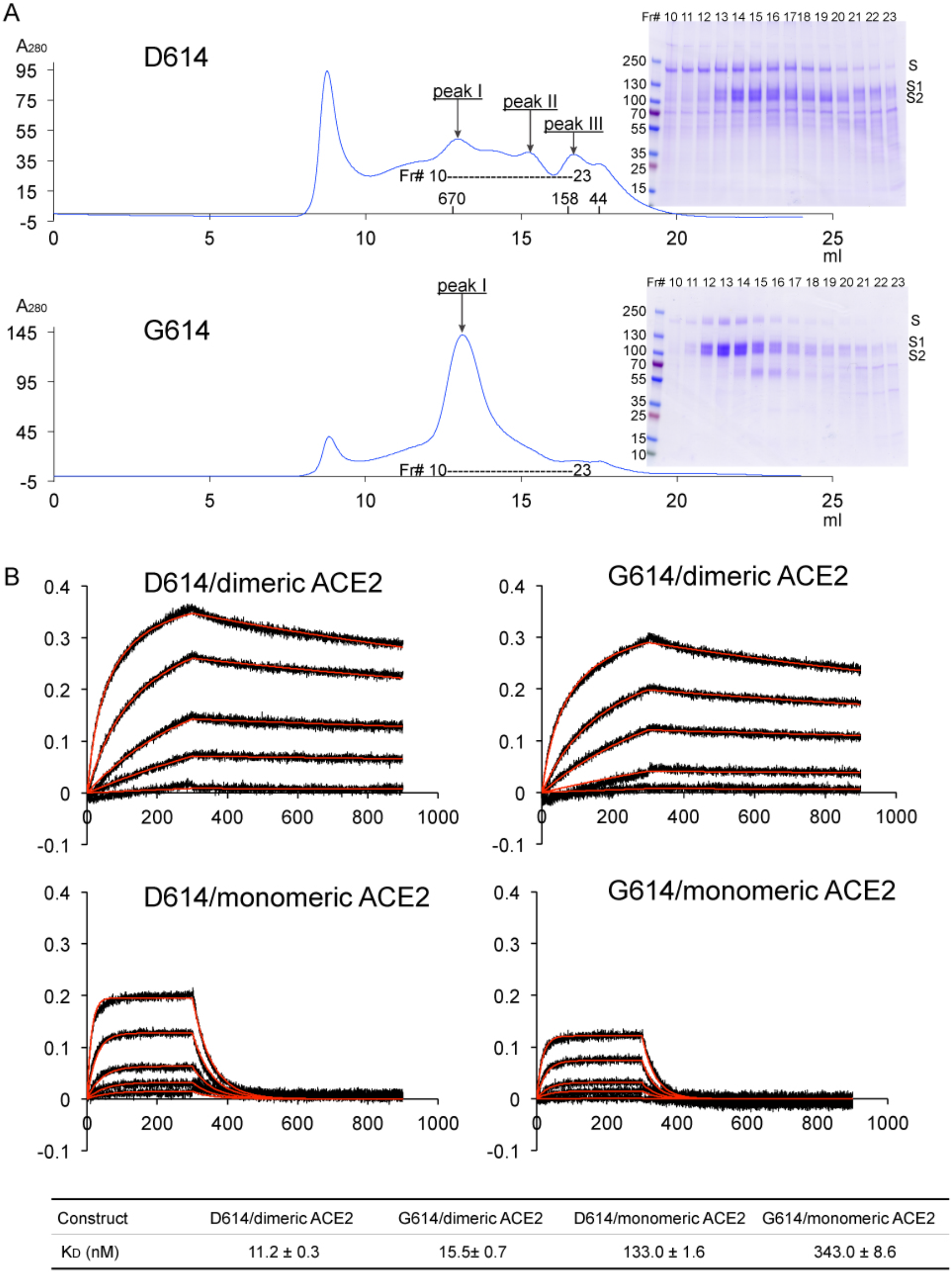
Characterization of the purified full-length SARS-CoV-2 S proteins. (A) The full-length SARS-CoV-2 S protein carrying either D614 or G614 was extracted and purified in detergent DDM, and further resolved by gel-filtration chromatography on a Superose 6 column. The molecular weight standards include thyoglobulin (670 kDa), γ-globulin (158 kDa) and ovalbumin (44 kDa). Inset, peak fractions were analyzed by Coomassie stained SDS-PAGE. Labeled bands are S, S1 and S2. Fr#, fraction number. (B) Binding analysis of fractions of Peak 1 in (A) with soluble ACE2 constructs by biolayer interferometry (BLI). The purified S proteins were immobilized to AR2G biosensors and dipped into the wells containing ACE2 at various concentrations (5.56-450 nM for monomeric ACE2, 2.78-225 nM for dimeric ACE2). Binding kinetics was evaluated using a 1:1 Langmuir binding model for the monomeric ACE2 and a bivalent model for dimeric ACE2. The sensorgrams are in black and the fits in red. Binding constants are also summarized here and in Table S1. All experiments were repeated at least twice with essentially identical results.

We measured by bio-layer interferometry (BLI) binding of the prefusion trimer fractions of the full-length proteins to recombinant soluble ACE2 (Fig. 1B). The S trimers bound more strongly to a dimeric ACE2 than to a monomeric ACE2, as expected. The G614 protein bound ACE2 less tightly than did the D614 protein, consistent with the measurements reported by others using soluble constructs^27^. This observation appears inconsistent with reports that the G614 trimer has a more exposed RBD than the D614 trimer^17,18,25,27^.

### Cryo-EM structures of the full-length S trimer with G614

We determined the cryo-EM structures of the full-length S trimer carrying G614 in both DDM and NP-40. Cryo-EM images were acquired on a Titan Krios electron microscope operated at 300 keV and equipped with a Gatan K3 direct electron detector. We used RELION^29^ for particle picking, two-dimensional (2D) classification, three dimensional (3D) classification and refinement. 3D classification identified three distinct classes each containing a similar number of particles. The three classes represent a closed, three RBD-down conformation, a one RBD-up conformation and an intermediate conformation with one RBD flipped up only halfway. All structures were refined to 3.1-3.5 Å resolution (Fig. S2-S5; Table S2). The same three classes were observed for the samples purified in both detergents, giving essentially identical maps of the corresponding classes after refinement (Fig. S6). These results demonstrate that detergent has little impact on the S structure at least in the visible regions of the ectodomain.

The overall structure of the full-length S protein with G614 in the closed, three RBD-down prefusion conformation is very similar to that of the D614 S trimer that we have published recently (Fig. 2; ref^15^). In the three RBD-down structure, the four domains in each S1, including NTD, RBD, CTD1 and CTD2, wrap around the three-fold axis of the trimer, protecting the prefusion S2. The furin cleavage site at the S1/S2 boundary remains disordered, making it difficult to determine whether this structure represents the uncleaved or cleaved trimer, although the preparation contains primarily the cleaved forms (Fig. 1A). The S2 fragment folds around a central three-stranded coiled coil that forms the most stable part of the structure with the strongest density in the entire S trimer; it is also the least variable region among all the known S trimer structures. The S2 structure is identical in the two structures of the G614 trimer with one RBD projecting upwards, either completely or partially (Figs. S7 and S8A). In the conformation with one RBD fully up, the two neighboring NTDs, including the one from the same protomer, shift away from the three-fold axis (Fig. S7). In the RBD-intermediate conformation, only the NTD from the adjacent protomer packing directly against the moving RBD shifted, suggesting that there is at least one local free-energy minimum along the pathway of the RBD upward movement.

**Figure 2.**
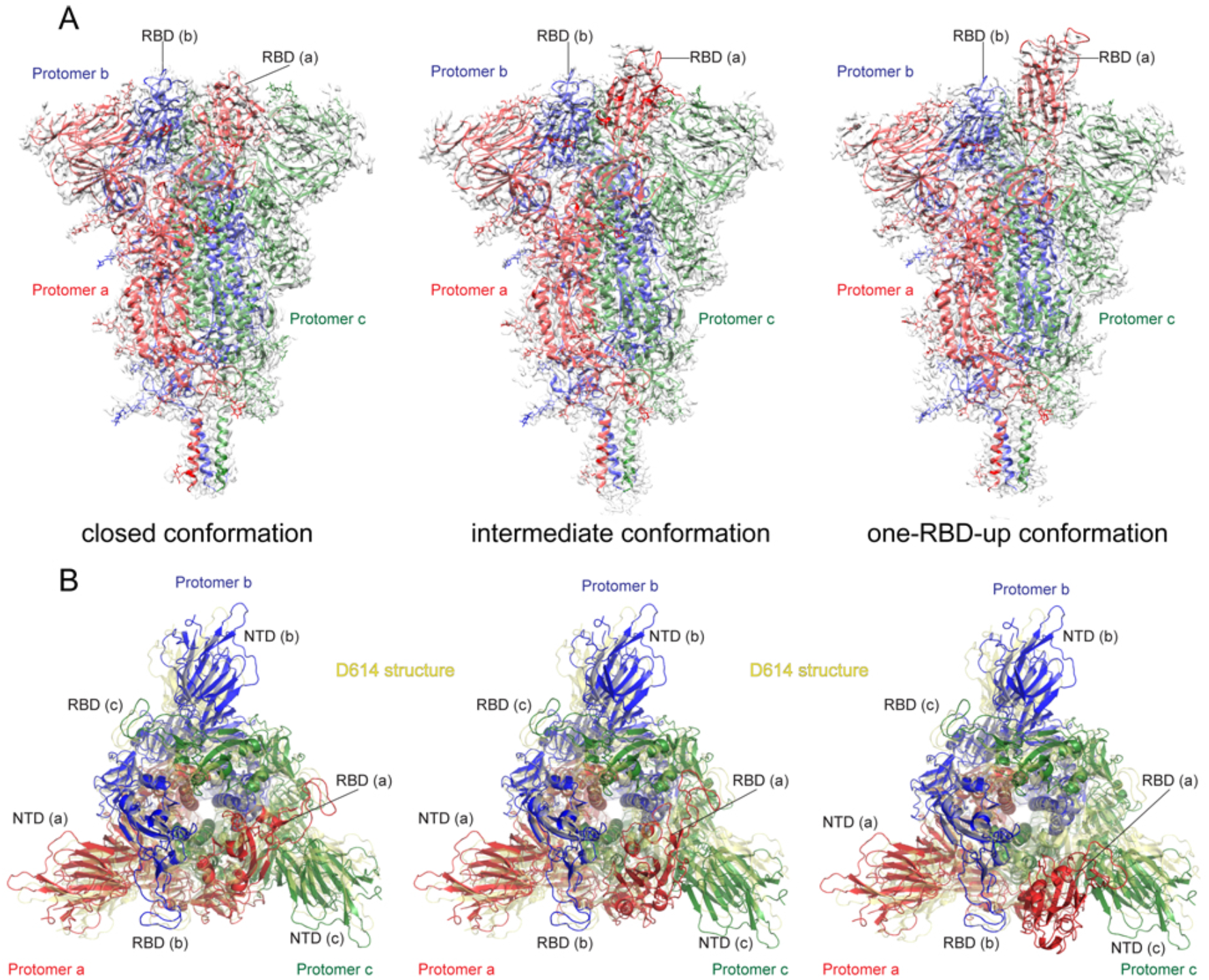
Cryo-EM structures of the full-length SARS-CoV-2 S protein carrying G614. (A) Three structures of the G614 S trimer, representing a closed, three RBD-down conformation, an RBD-intermediate conformation and a one RBD-up conformation, were modeled based on corresponding cryo-EM density maps at 3.1-3.5Å resolution. Three protomers (a, b, c) are colored in red, blue and green, respectively. RBD locations are indicated. (B) Top views of superposition of three structures of the G614 S in (A) in ribbon representation with the structure of the prefusion trimer of the D614 S (PDB ID: 6XR8), shown in yellow. NTD and RBD of each protomer are indicated. Side views of the superposition are shown in Fig. S8.

The D614G substitution eliminates a salt bridge between residue 614 in CTD2 of one subunit and residue 854 in the FPPR of the adjacent subunit, probably destabilizing the latter^15^. Nonetheless, the FPPR in the three RBD-down conformation of the G614 trimer is structured, although the density in the regions is slightly weaker than in the D614 map (Fig. S8B). A major difference between the G614 and D614 trimer structures is that a ~20-residue segment (620-640) in the CTD2, largely disordered in the D614 trimer, has become structured in the G614 trimer. There is no density for the C-terminal segments, including HR2, TM and CT, consistent with the flexibility near residue Pro1162 found in cryo-ET studies^17,18^.

### Structural consequences of the D614G substitution

To examine the structural changes resulted from the D614G substitution, we superposed the structures of the G614 trimer onto the D614 trimer in the closed conformation aligning them by the invariant S2 (Fig. 2B). A shift by a clockwise, outward rotation of all three S1 subunits, relative to the D614 structure, is evident even for the G614 trimer in the closed conformation. A similar shift was also observed in the RBD-intermediate and RBD-up G614 structures. Thus, the D614G substitution has led to a slightly more open conformation than that of the D614 trimer, even when all three RBDs are down. The D614G change has apparently also rigidified a neighboring segment of CTD2, residues 620-640, which we designate the “630 loop”. This loop inserts into a gap, slightly wider in the G614 than in the D614 trimer, between the NTD and CTD1 of the same protomer (Figs. 3 and 4). The 630 loop is disordered in the closed D614 trimer, because the gap narrows enough to exclude it. The close D614 trimer thus has three ordered FPPRs and three disordered 630 loops, while in the closed G614 trimer structures reported here, the three 630 loops and three FPPRs in the closed conformation are all structured. In the two conformers with one partly or fully open RBD, the two segments are disordered in the RBD-shifted subunit, and their central parts have difficult-to-model density in one other subunit. The third pair appears well ordered throughout. Thus, opening of the RBD in the full-length, G614 trimer correlates with a displacement of the 630 loop and the FPPR away from their positions in the D614 variant, making them too mobile to be detected in our cryo-EM reconstructions.

**Figure 3.**
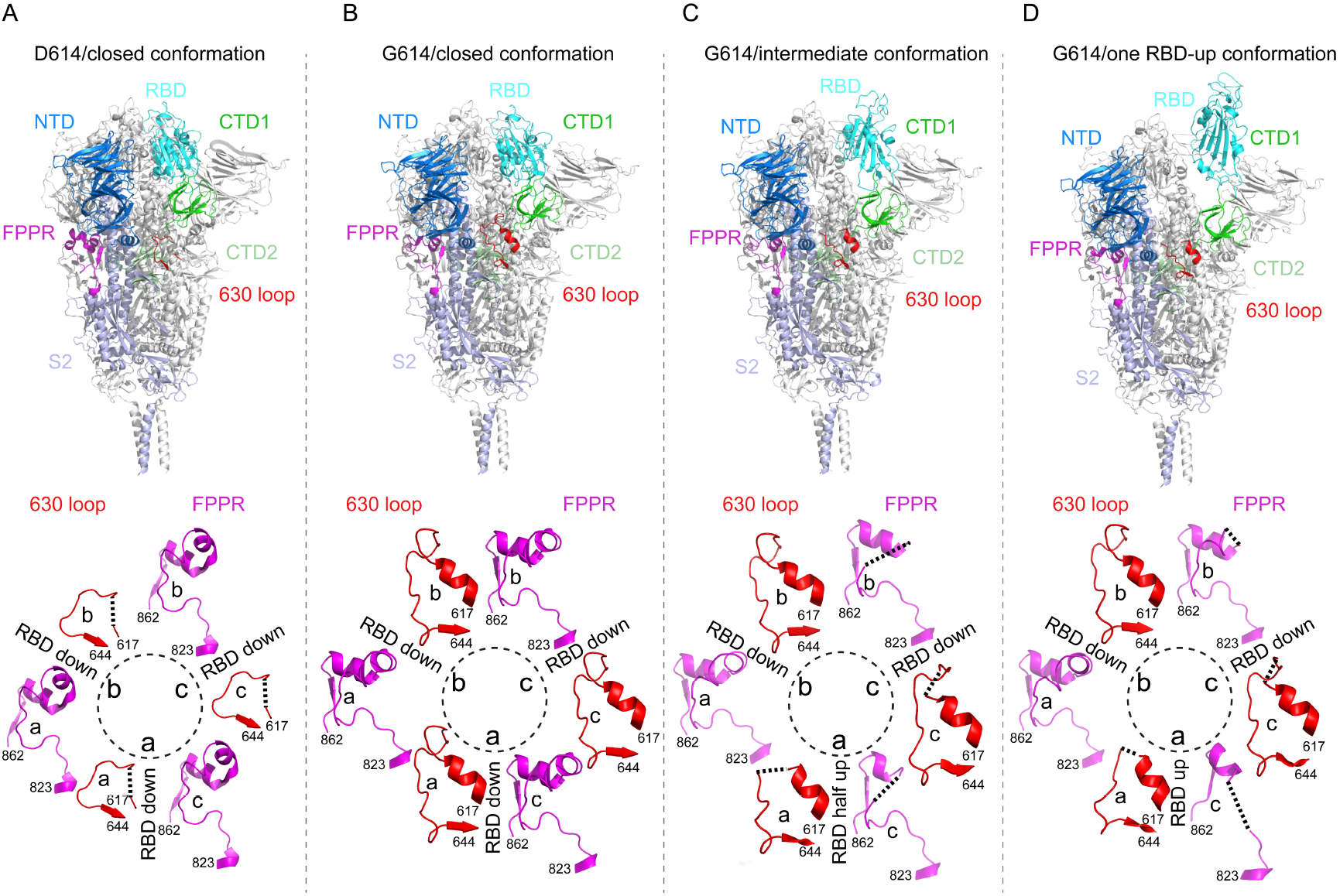
Cryo-EM structures of the full-length SARS-CoV-2 S protein carrying G614. (A) Top, the structure of the closed, three RBD-down conformation of the D614 S trimer is shown in ribbon diagram with one protomer colored as NTD in blue, RBD in cyan, CTD1 in green, CTD2 in light green, S2 in light blue, the 630 loop in red and the FPPR in magenta. Bottom, structures of three segments (residues 617-644) containing the 630 loop in red and three segments (residues 823-862) containing the FPPR in magenta from all three protomers (a, b and c) are shown. Position of each RBD is indicated. (B-D) Structures of the G614 trimer in the closed, three RBD-down conformation, the RBD-intermediate conformation and the one RBD-up conformation, respectively, are shown as in (A). Dash lines indicate gaps.

**Figure 4.**
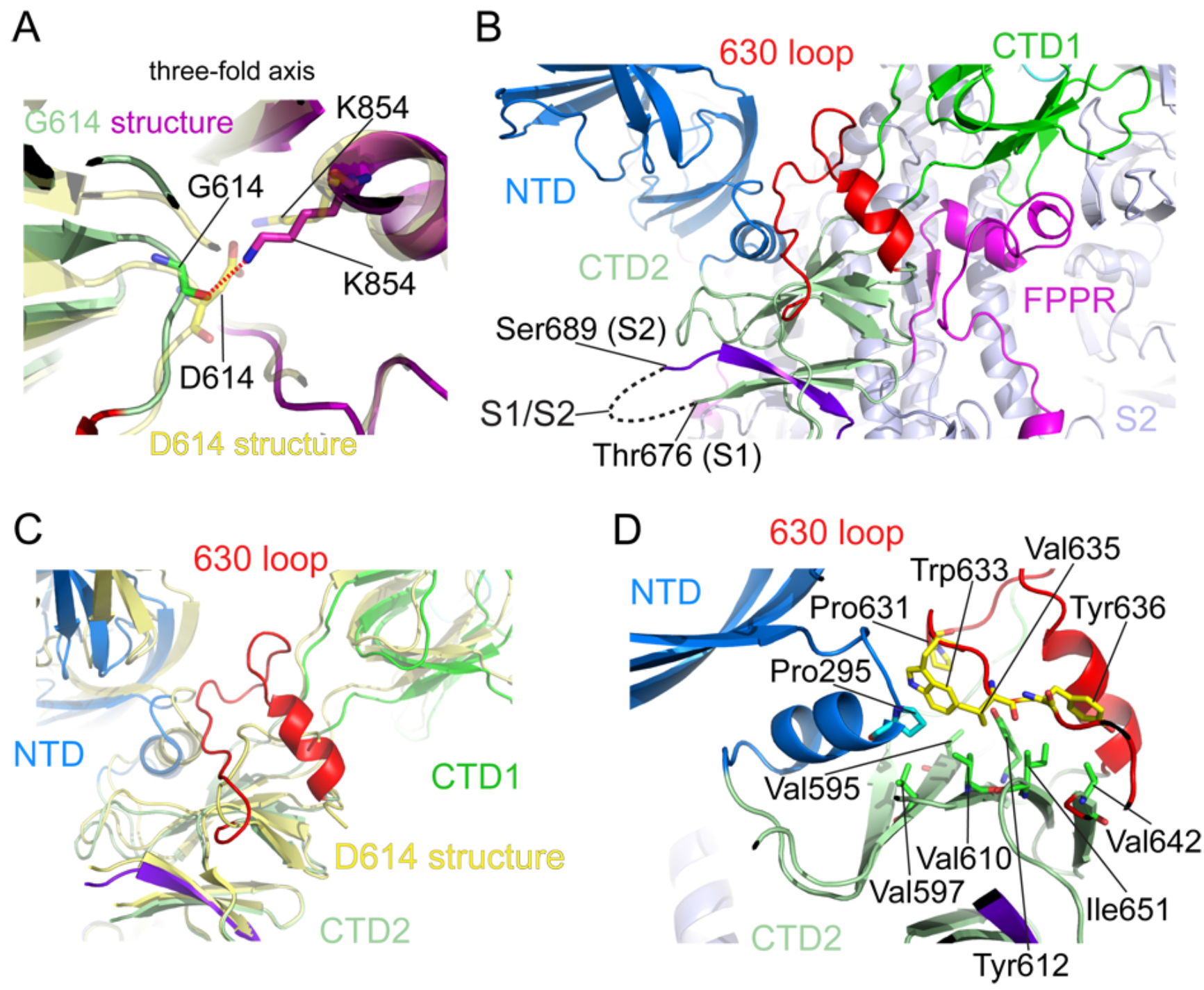
Close-up views of the D614G substitution. (A) A close-up view of the region near the residue 614 with superposition of the G614 trimer structure in green (CTD2) and magenta (FPPR) and the D614 trimer in yellow, both in the closed prefusion conformation. Residues G614, D614 and two K854 from both structures are shown in stick model. The direction of the three-fold axis of the trimer is indicated. The hydrogen bond between G614 and K854 is indicated by a red dash line. (B) Location of the 630 loop in the S trimer. The 630 loop is highlighted in red, NTD in blue, CTD1 in green, CTD2 in light green, S2 in light blue, and the FPPR from a neighboring protomer in magenta. The S1/S2 boundary and the nearest ordered residues Thr676 from S1 and S689 from S2 are all indicated. A strand from the N-terminal end of S2, packed in the CTD2, is highlighted in purple. (C) A view showing that the 630 loop wedges between the NTD and the CTD1 and pushed them apart. (D) Packing of the 630 loop against the hydrophobic surface formed by residues Val595, Val597, Val610 Tyr 612, Val642 and Ile 651 from the CTD2 and Pro295 from the NTD. Residues Pro631, Trp633 and Val635 from the 630 loop contribute to this interaction.

In the immediate vicinity of residue 614, the change from Asp to Gly did not cause any large local structural rearrangements except for loss of the D614-K854 salt bridge we had previously predicted^15^, and a small shift of residue 614 towards the three-fold axis (Fig. 4A). The position of the FPPR and the conformation of K854 allow a hydrogen bond between K854Nη and the main-chain carbonyl of G614, perhaps accounting for the subtlety of the structural difference. The loss of the salt bridge involving D614, at least partially compensated by the new hydrogen bond between G614 and K854, was apparently not sufficient to destabilize the packing of the FPPR against the rest of the trimer, but it did weaken the FPPR density, especially between residues 842-846. The 630 loop, which packs directly against the NTD, CTD1 and CTD2 of the same protomer, lies close to the S1/S2 boundary of the same protomer and the FPPR of an adjacent protomer (Fig. 4B). The loop inserts between the NTD and CTD1 (Fig. 4C), probably requiring the shifts illustrated in Fig. 2B. This wedge-like loop may also help secure the NTD and CTDs and enhance G614 S trimer stability.

CTD2 is formed by two stacked, four-strand β-sheets, with a fifth strand in one sheet contributed by the connector between the NTD and RBD. In the other sheet, an interstrand loop contains the S1/S2 cleavage site, and thus one strand is the N-terminal segment of S2 (Fig. 4B). In the G614 trimer, one side of the 630 loop packs along a long hydrophobic surface, largely solvent-exposed in the D614 trimer, formed by residues on the “upward” facing surface of the CTD2 along with Pro295 from the NTD (Fig. 4D). Pro631, Trp633 and Val635 of the 630 loop appear to contribute to this interaction. Since S1 dissociation from S2 likely requires destabilization of the CTD2 to free the β-strand from the N-terminal end of S2, an ordered 630 loop that completes folding of the CTD2 by closing off an exposed, hydrophobic surface may retard S1 shedding, thereby enhancing the stability of a cleaved S trimer.

We note that although S1 in the G614 trimer moves outwards from its position in the D614 trimer, the extent of the shift is still appreciably smaller than the shift seen in soluble S trimers stabilized by a trimerization foldon tag and two proline mutations (Fig. S9). The comparison suggests that the soluble trimer may not completely mimic all the physiologically relevant conformations of the S trimer.

## Discussion

### Structural basis of increased stability and infectivity of the G614 spike

The structures described here allow us to propose an explanation why the virus carrying the D614G substitution, with a more stable S trimer, is more infectious than the original strain. We consider the schematic free energy landscape of the S-protein conformational distribution in Fig. 5. In the D614 trimer, the kinetic barrier for transition from the RBD-down conformation, clamped by the FPPR only, to the RBD-up conformation, is relatively low, and so is for the barrier for the subsequent irreversible transition to the postfusion S2 and dissociated S1. The second transition can occur even in the absence of ACE2, as seen on the surfaces of pseudotyped viruses and inactivated SARS-CoV-2 that carry D614^20,26^, as well as our recent study^15^. Since the RBD-up conformation could either revert to the closed state or dissociate irreversibly to allow transition to the postfusion state, it is not surprising that very few such particles were captured in our previous cryo-EM study or in the cryo-ET study of the inactivated virus using the D614 variant^15,20^. Instead, the closed prefusion conformation and the postfusion conformation were the principal species observed. In the G614 trimer, the RBD-down conformation is reinforced not only by the FPPR but also by the newly identified 630 loop, substantially raising the barrier for the closed-to-open transition of the RBD. Local free energy minima could trap some RBD intermediate conformations, perhaps accounting for the RBD-intermediate captured here.

**Figure 5.**
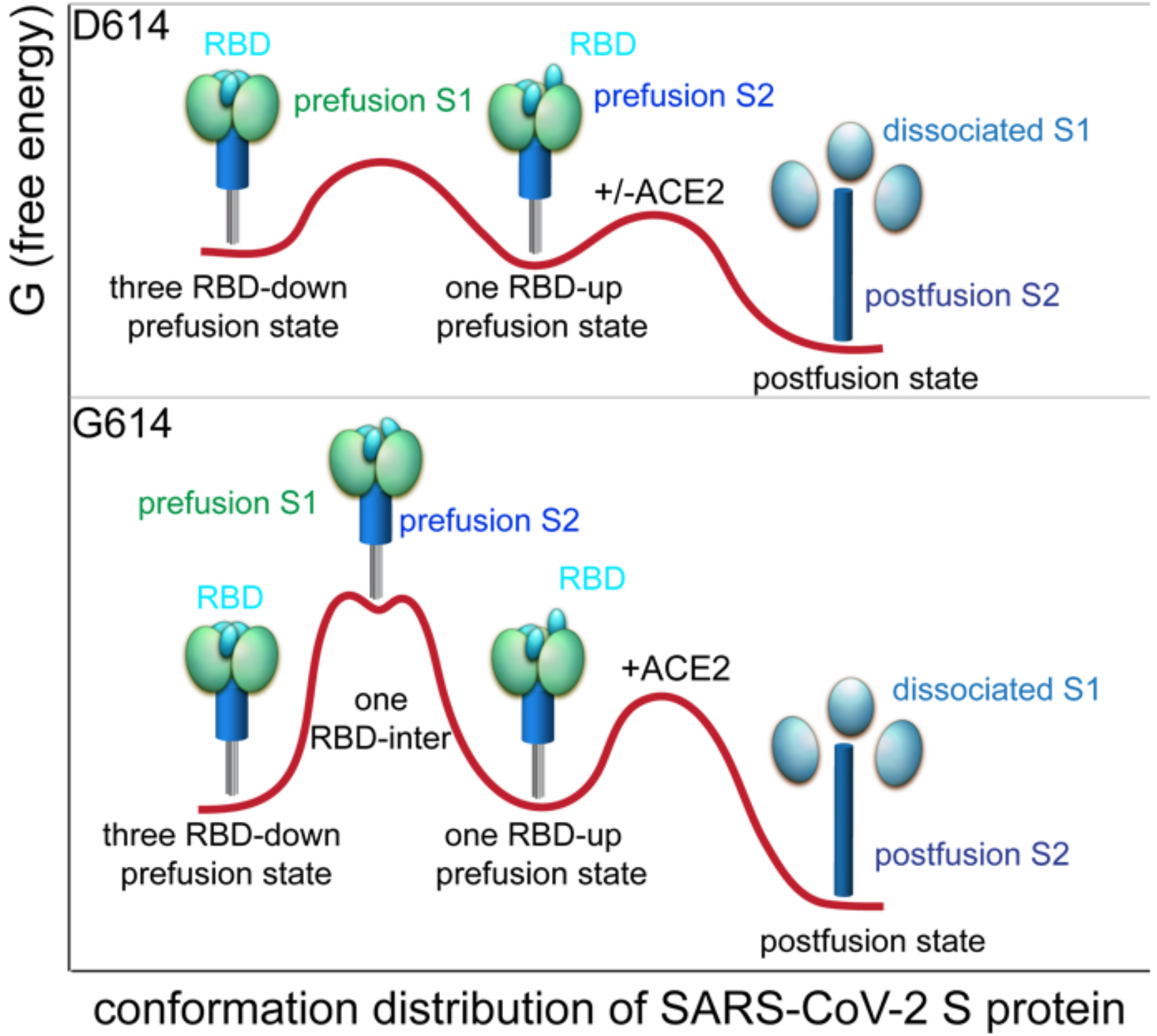
Proposed conformation distribution of the SARS-CoV-2 S trimer. Free energy (G) landscapes are proposed for the D614 S trimer that samples the closed, one RBD-up and postfusion conformations and for the G614 S trimer that samples the closed, RBD-intermediate, one RBD-up and postfusion conformations.

Our interpretation of the structural differences is also consistent with the spike conformational distribution on the virions in cryo-ET studies of chemically inactivated SARS-CoV-2. The D614 preparation contains primarily postfusion S2 spikes^20^. One study of a G614 virus that had lost the furin cleavage site showed almost no postfusion spikes, and a 50:50 distribution of prefusion spikes between fully closed and one RBD-up^17^; and another showed 3% postfusion spikes and 97% in the prefusion form (~31%, fully closed; ~55%, one RBD-up; ~14% two RBD-up)^18^. The structured 630 loop in the G614 trimer not only reinforces the packing among three protomers by inserting between the NTD and CTD1 of the same protomer but also seals a hydrophobic surface of the CTD2. The latter interaction probably stabilizes the domain and thus inhibits release of the N-terminal segment of S2, effectively blocking S1 dissociation. This property can account for the paucity of postfusion spikes on the G614 variant.

Our data can also explain why the G614 trimer binds somewhat less tightly to ACE2 than does the D614 trimer, despite the greater proportion of well-exposed RBDs, both as seen here and as reported by others^17,18,25,27^. The higher kinetic barrier for an upward shift in the RBD than that in the D614 trimer would create a significant hurdle for the rest of RBDs in the same G614 trimer or the closed G614 trimers to adopt an ACE2-accessible state. We note that the second binding event with the dimeric ACE2 has a slower on-rate and also a slower off-rate for the G614 than the D614 trimer (Table S1).It is therefore not surprising that the G614 trimer preparation showed weaker binding to both monomeric and dimeric ACE2 than did a D614 S preparation. We suggest that the enhanced infectivity of the G614 virus largely results from the increased stability of the S trimer, rather than the better exposed RBDs. Indeed, if the virus that passed from bats to humans or to an intermediate vector contained D614 (also present in the bat coronavirus BatCoV RaTG13^1^), then it could have gained fitness in the new host by acquiring changes such as G614 for greater stability and infectivity than the parental form.

### Membrane fusion mechanism

We previously hypothesized that the FPPR might modulate the fusogenic structural rearrangements of S protein, as it retains the RBDs in the down conformation but moves out of its position when the RBD flips up. The current study has shown that additional structures, CTD2 and the 630 loop within it, are probably key components of the S fusion machinery. If ACE2 captures the RBD-up conformation, expelling both the 630 loop and the FPPR from their positions in the closed S trimer conformation, the FPPR shift may help expose the S2’ site near the fusion peptide for proteolytic cleavage, while departure of the 630 loop from the hydrophobic surface of the CTD2 can destabilize this domain and free the N-terminal segment of S2 to dissociate from S1, if the furin site has already been cleaved, and release S1 altogether. Dissociation of S1 would then initiate a cascade of refolding events in the metastable prefusion S2, allowing the fusogenic transition to a stable postfusion structure. We note that this model is very similar to that proposed for membrane fusion catalyzed by HIV envelope protein, in which gp120 dissociation triggers refolding of gp41 to complete the fusion process^30^.

### Implications for vaccine development

The SARS-CoV-2 S protein is the centerpiece of almost all the first-generation vaccines, being developed, which began at early stages in the pandemic and used the D614 sequence. We have previously suggested that the inactivated-virus vaccines might have too many postfusion spikes and induce mainly non-neutralizing antibodies^15^. Indeed, these vaccine candidates induced the lowest level of neutralizing antibody responses; other S constructs containing stabilization modifications to prevent conformational changes have given much stronger responses^31^, but their protective efficacy still needs to be evaluated in phase 3 clinical trials. The G614 S trimer is naturally constrained in a prefusion state that presents both the RBD-down and RBD-up conformations with great stability. It is therefore likely to be a superior immunogen, whether in a protein or nucleic acid form, for eliciting protective neutralizing antibody responses, which appear largely to target the RBD and NTD^32,33^.

## Acknowledgments

We thank the SBGrid team for technical support, K. Arnett for support and advice on the BLI experiments, and S. Harrison, M. Liao, A. Carfi and D. Barouch for critical reading of the manuscript. EM data were collected at the Harvard Cryo-EM Center for Structural Biology of Harvard Medical School. This work was supported by NIH grants AI147884 (to B.C.), AI147884-01A1S1 (to B.C), AI141002 (to B.C.), AI127193 (to B.C. and James Chou), a COVID19 Award by Massachusetts Consortium on Pathogen Readiness (MassCPR; to B.C.), as well as a Fast grant by Emergent Ventures (to B.C.).

## Author Contribution

B.C. J.Z., Y.C. and T.X. conceived the project. Y.C. and H.P. expressed and purified the full-length S proteins. T.X. expressed and purified soluble ACE2 constructs with help from H.P.. T.X. performed BLI and cell-cell fusion experiments. J.Z. prepared cryo grids and performed EM data collection with contributions from S.M.S. and R.M.W.. J.Z. processed the cryo-EM data, built and refined the atomic models. J.L. created the G614 expression construct. S.R.V. contributed to cell culture and protein production. P.S. provided computational support. All authors analyzed the data. B.C., J.Z., Y.C. and T.X. wrote the manuscript with input from all other authors.

## Methods

### Expression constructs

Genes of a full-length spike (S) protein (residue 1-1273) of SARS-CoV-2 (D614) and the ectodomain of human angiotensin converting enzyme 2 (ACE2; residue 1-615) were synthesized by GenScript (Piscataway, NJ). The S gene was fused with a C-terminal twin Strep tag [(GGGGS)2WSHPQFEK(GGGGS)2WSHPQFEK)] and cloned into a mammalian cell expression vector pCMV-IRES-puro (Codex BioSolutions, Inc, Gaithersburg, MD). The D614G substation in the S gene was generated by site-specific mutagenesis and confirmed by DNA sequencing. The ACE2 gene added a C-terminal 6xhistidine tag and cloned into the same expression vector. The dimeric ACE2 construct was kindly provided by Dr. Michael Farzan at Scripps Research Institute.

### Expression and purification of recombinant proteins

Expression and purification of the full-length S protein carrying G614 were carried out as previously described^15^. Briefly, expi293F cells were transiently transfected with the S protein expression construct. To purify the S protein, the transfected cells were lysed in a buffer containing Buffer A (100 mM Tris-HCl, pH 8.0, 150 mM NaCl, 1 mM EDTA) and 1% (w/v) n-dodecyl-β-D-maltopyranoside (DDM) (Anatrace, Inc. Maumee, OH), EDTA-free complete protease inhibitor cocktail (Roche, Basel, Switzerland), and incubated at 4°C for one hour. After a clarifying spin, the supernatant was loaded on a strep-tactin column equilibrated with the lysis buffer. The column was then washed with 50 column volumes of Buffer A and 0.3% DDM, followed by additional washes with 50 column volumes of Buffer A and 0.1% DDM, and with 50 column volumes of Buffer A and 0.02% DDM. The S protein was eluted by Buffer A containing 0.02% DDM and 5 mM desthiobiotin. The protein was further purified by gel filtration chromatography on a Superose 6 10/300 column (GE Healthcare) in a buffer containing 25 mM Tris-HCl, pH 7.5, 150 mM NaCl, 0.02% DDM. Similar purification was also performed using detergent NP-40 instead, as described previously for the D614 S trimer^15^.

Expi293F cells transfected with monomeric ACE2 or dimeric ACE2 expression construct were grown in 250 ml roller bottles with DMEM containing 10% FBS. The supernatant of the cell culture was collected by centrifugation at 2,524 xg for 30 minutes. The monomeric ACE2 was purified by affinity chromatography using Ni-NTA agarose (Qiagen, Hilden, Germany), followed by gel filtration chromatography, as described previously^34,35^. The peak fractions were pooled and concentrated to 10 mg/ml using a 30 kDa MWCO Millipore filter (MilliporeSigma, Burlington, MA). The supernatant of dimeric ACE2 was loaded to a column packed with GammaBind Plus Sepharose beads (GE Healthcare). The column was washed with PBS. The protein was eluted using 100 mM glycine (pH 2.5) and neutralized immediately with 2 M Tris-HCl (pH 8.0). The eluted protein was further purified by gel filtration chromatography on a Superdex 200 Increase 10/300 GL column. The peak fractions were pooled and concentrated to 5 mg/ml using a 50kDa MWCO Millipore filter.

### Binding assay by bio-layer interferometry (BLI)

Binding of monomeric or dimeric ACE2 to the full-length Spike protein was measured using an Octet RED384 system (ForteBio, Fremont, CA). Each ACE2 protein was diluted using the running buffer (PBS, 0.02% Tween 20, 1 mg/ml BSA) and transferred to a 96-well plate. The full-length S protein was immobilized to Amine Reactive 2^nd^ Generation (AR2G) biosensors (ForteBio), following a protocol recommended by the manufacturer. After equilibrating in the running buffer for 5 minutes, the sensors with immobilized Spike protein were dipped in the wells containing the ACE2 protein at various concentrations (5.56-450 nM for monomeric ACE2; 2.78-225 nM for dimeric ACE2) for 5 minutes to measure the association rate. The sensors were then dipped in the running buffer for 10 minutes to determine the dissociation rate. Control sensors with no Spike protein were also dipped in the ACE2 solutions and the running buffer as references. Recorded sensorgrams with background subtracted from the references were analyzed using the software Octet Data Analysis HT Version 11.1 (ForteBio). The curves for monomeric ACE2 were fit to a 1:1 binding model, while those for dimeric ACE2 were fit to a bivalent binding model.

### Cell-cell fusion assay

The cell-cell fusion assay, based on the α-complementation of E. coli β-galactosidase, was conducted to quantify the fusion activity mediated by SARS-CoV2 S protein, as described^15^. Briefly, various amount of the full-length SARS-CoV2 (614D or 614G) S construct (0.025-10 μg) and the α fragment of E. coli β-galactosidase construct (10 μg), or the full-length ACE2 construct (10 μg) together with the ω fragment of E. coli β-galactosidase construct (10 μg), were transfected to HEK293T cells using Polyethylenimine (PEI) (80 μg). After a 24-hour incubation at 37°C, the cells were detached using DPBS buffer with 5mM EDTA and resuspended in complete DMEM medium. 50 μl S-expressing cells (1.0×10^6^ cells/ml) were mixed with 50 μl ACE2-expressing cells (1.0×10^6^ cells/ml) to allow the cell-cell fusion proceed at 37 °C for 2 hours. Cell-cell fusion activity was quantified using a chemiluminescent assay system, Gal-Screen (Applied Biosystems, Foster City, CA), following the standard protocol recommended by the manufacturer. The substrate was added to the mixture of the cells and allowed to react for 90 minutes in dark at room temperature. The luminescence signal was recorded with a Synergy Neo plate reader (Biotek).

For the inhibition assay, the S-expressing cells were incubated with trimeric ACE2 variant, ACE2_615_-foldon T27W^28^, at varied concentrations (6.25-200 μg/ml) for 1 hour at 37°C. After the incubation, the ACE2-expressing cells were added to the mixture, followed with a 2-hour incubation at 37 °C. The fusion activity was quantified using the Gal-Screen system as mentioned above.

### Cryo-EM sample preparation and data collection

To prepare cryo grids, 3.5 μl of the freshly purified G614 sample in NP-40 at ~0.3 mg/ml was applied to a 1.2/1.3 Quantifoil grid with continuous carbon support (Quantifoil Micro Tools GmbH), which had been glow discharged with a PELCO easiGlow™ Glow Discharge Cleaning system (Ted Pella, Inc.) for 60 s at 15 mA. For the G614 sample in DDM, 3.5 μl of the peak fraction from gel filtration chromatography at ~1.0 mg/ml was also applied to the glow discharged 1.2/1.3 Quantifoil grids (Quantifoil Micro Tools Gmb). Grids were immediately plunge-frozen in liquid ethane using a Vitrobot Mark IV (ThermoFisher Scientific), and excess protein was blotted away by using grade 595 filter paper (Ted Pella, Inc.) with a blotting time of 4 s, a blotting force of -12 at 4□ in 100% humidity. The grids were first screened for ice thickness and particle distribution using a Talos Arctica transmission electron microscope (ThermoFisher Scientific), operated at 200 keV and equipped with a K3 direct electron detector (Gatan). For data collection, images were acquired with selected grids using a Titan Krios transmission electron microscope (ThermoFisher Scientific) operated at 300 keV and equipped with a BioQuantum GIF/K3 direct electron detector. Automated data collection was carried out using SerialEM version65 ^36^ at a nominal magnification of 105,000× and the K3 detector in counting mode (calibrated pixel size, 0.825 Å) at an exposure rate of ~14.7 electrons per physical pixel per second for the two different detergent samples (14.77 and 14.68, respectively). Each movie had a total accumulated electron exposure of ~50 e/Å^2^, fractionated in 50 frames (50 ms per frame). Datasets were acquired using a defocus range of 1.5-2.7 μm for the NP-40 sample and 1.0-2.5 μm for the DDM sample.

### Image processing and 3D reconstructions

Drift correction for cryo-EM images was performed using MotionCor2^37^, and contrast transfer function (CTF) was estimated by CTFFIND4^38^ using motion-corrected sums without dose-weighting. Motion corrected sums with dose-weighting were used for all other image processing. RELION3.0.8 was used for particle picking, 2D classification, 3D classification and refinement procedure. Approximately 3,000 particles were manually picked for each protein sample and subjected to 2D classification to generate the templates for automatic particle picking. For the NP-40 sample, after manual inspection of auto-picked particles, a total of 3,640,242 particles were extracted from 11,577 images. The selected particles were subjected to 2D classification, giving a total of 2,657,624 good particles. A low-resolution negative-stain reconstruction of the sample was low-pass filtered to 30Å resolution and used as an initial model for 3D classification with C3 symmetry. One major class showed clear structural features were subjected to another round of 3D classification with C1 symmetry, giving one major class. Third round of 3D classification with C1 symmetry and further local angular search produced three major classes, representing the closed, three RBD-down conformation, the one RBD-up conformation and the RBD-intermediate conformation, respectively. The three classes were then subjected to 3D auto-refinement, followed by particle polishing and signal-subtraction classification focused on the apex region of the S trimer. The three classes containing 63,558, 86,353 and 93,426 particles were subjected to 3D refinement with C1 (RBD-intermediate and RBD-up) and C3 (closed) symmetry using an overall mask, resulting in three final reconstructions at 3.3Å, 3.5Å and 3.1Å resolutions, respectively. For the DDM sample, a total of 5,652,781 particles were extracted from 11,092 images by reference-based auto-picking. Two rounds of 2D classification were performed, giving 3,207,721 good particles. These particles were subjected to two rounds of 3D classification with C3 symmetry, yielding one major class with clear structural features. The refined map from the first dataset was low-pass filtered to 30Å resolution and used as the initial model for 3D classification. A third round of 3D classification with C1 symmetry and further local angular search were performed, generating three major classes, which are similar to those from the NP-40 sample. CTF refinement and Bayesian polishing were performed for these classes. The RBD-up class containing 55,537 particles gave a 3D reconstruction at 3.2Å resolution after 3D auto-refine with C1 symmetry and overall mask; the RBD-intermediate class with 63,040 particles and the closed conformation class with 55,096 particles were subjected to the same refinement strategy, yielding 3D maps with resolution at 3.2Å and 3.4Å. The best map from each class was used for model building.

Reported resolutions are based on the gold-standard Fourier shell correlation (FSC) using the 0.143 criterion. All density maps were corrected from the modulation transfer function of the K3 detector and then sharpened by applying a temperature factor that was estimated using post-processing in RELION. Local resolution was determined using RELION with half-reconstructions as input maps.

### Model building

The initial templates for model building used the stabilized SARS-CoV-2 S ectodomain trimer structure (PDB ID 6XR8) for the prefusion conformation. Several rounds of manual building were performed in Coot^39^. The model was then refined in Phenix^40^ against the 3.1Å (closed), 3.2Å (RBD-intermediate) and 3.5Å (RBD-up) cryo-EM maps. Iteratively, refinement was performed in both Phenix (real space refinement) and ISOLDE^41^, and the Phenix refinement strategy included minimization_global, local_grid_search, and adp, with rotamer, Ramachandran, and reference-model restraints, using 6XR8 as the reference model. The refinement statistics are summarized in Table S2. Structural biology applications used in this project were compiled and configured by SBGrid^42^.

