## Supplemental Figures and Tables for "Structural impact on SARS-CoV-2 spike protein by D614G substitution"

### Supplementary materials

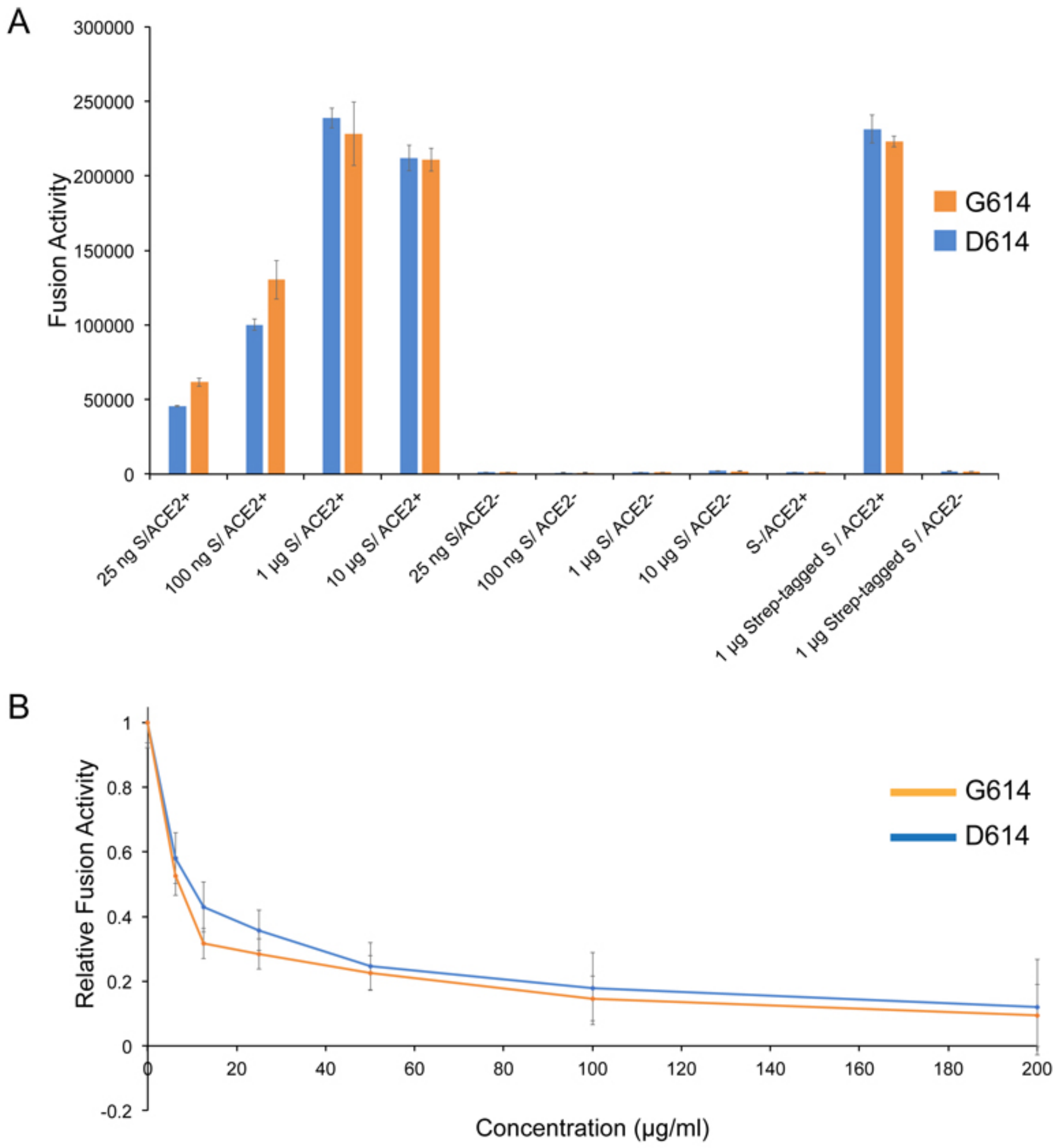


**Figure S1. Cell-cell fusion mediated by SARS-CoV-2 S protein carrying either D614 or G614 and its inhibition.** (A) HEK293T cells transfected with either the untagged or strep-tagged full length S protein expression plasmids containing D614 or G614 were fused with ACE2-expressing cells. Cell-cell fusion led to reconstitution of α and ω fragments of β-galactosidase yielding an active enzyme and thus the fusion activity was quantified by a chemiluminescent assay. No ACE2 and no S were negative controls. (B) Inhibition of the cell-cell fusion by a designed ACE2-based inhibitor $ACE2_{615}$-foldon-T27W as described[28].

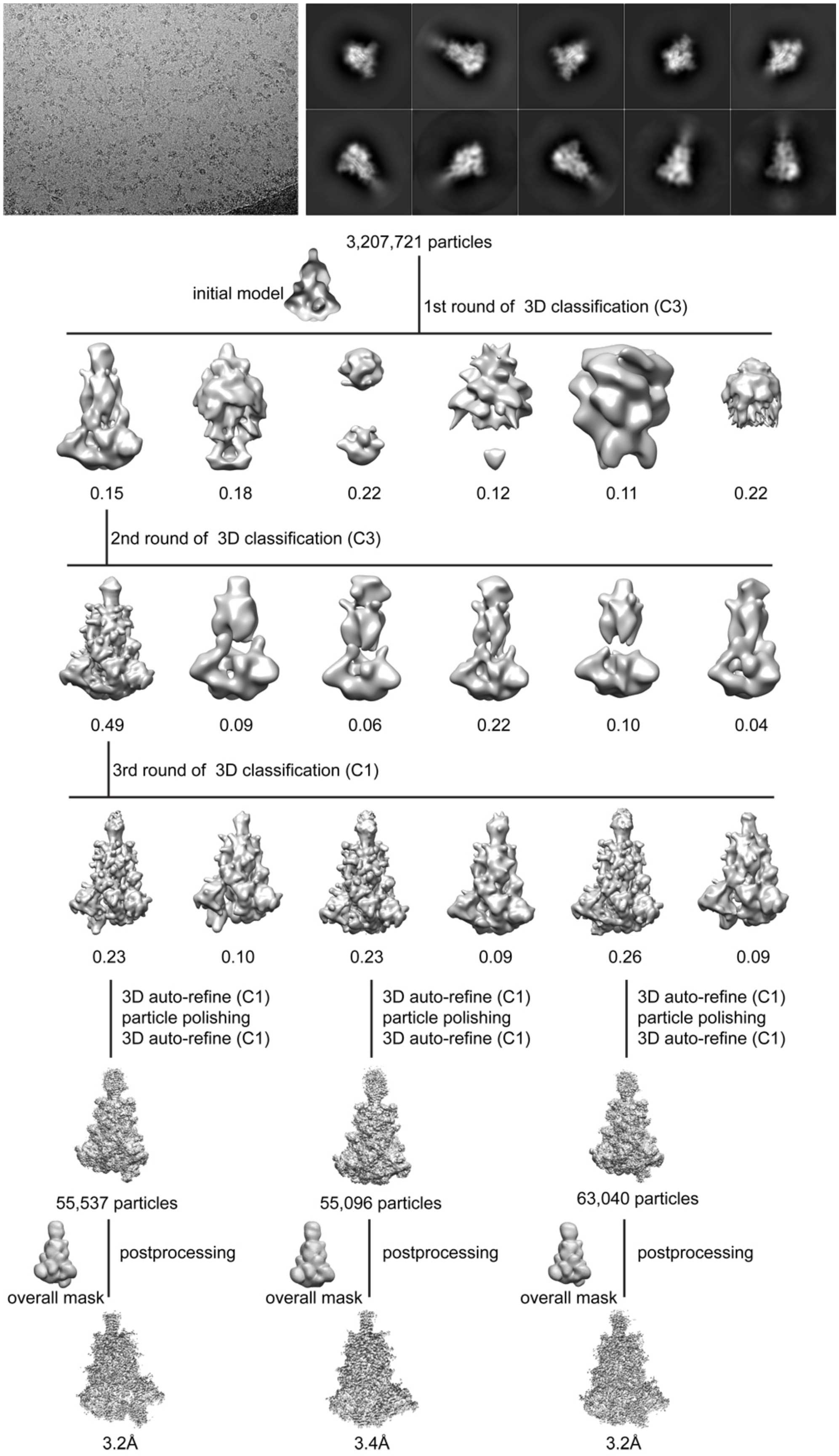
3,207,721 particles
initial model
1st round of 3D classification (C3)
0.15
0.18
0.22
0.12
0.11
0.22
2nd round of 3D classification (C3)
0.49
0.09
0.06
0.22
0.10
0.04
3rd round of 3D classification (C1)
0.23
0.10
0.23
0.09
0.26
0.09
3D auto-refine (C1)
particle polishing
3D auto-refine (C1)
3D auto-refine (C1)
particle polishing
3D auto-refine (C1)
3D auto-refine (C1)
particle polishing
3D auto-refine (C1)
55,537 particles
55,096 particles
63,040 particles
postprocessing
overall mask
postprocessing
overall mask
postprocessing
overall mask
3.2Å
3.4Å
3.2Å

**Figure S2. Cryo-EM analysis of the G614 S trimer in detergent DDM.** Top, representative micrograph, and 2D averages (box dimension: 396Å) of the cryo-EM particle images of the G614 S trimer in detergent DDM. Bottom, data processing workflow for structure determination.

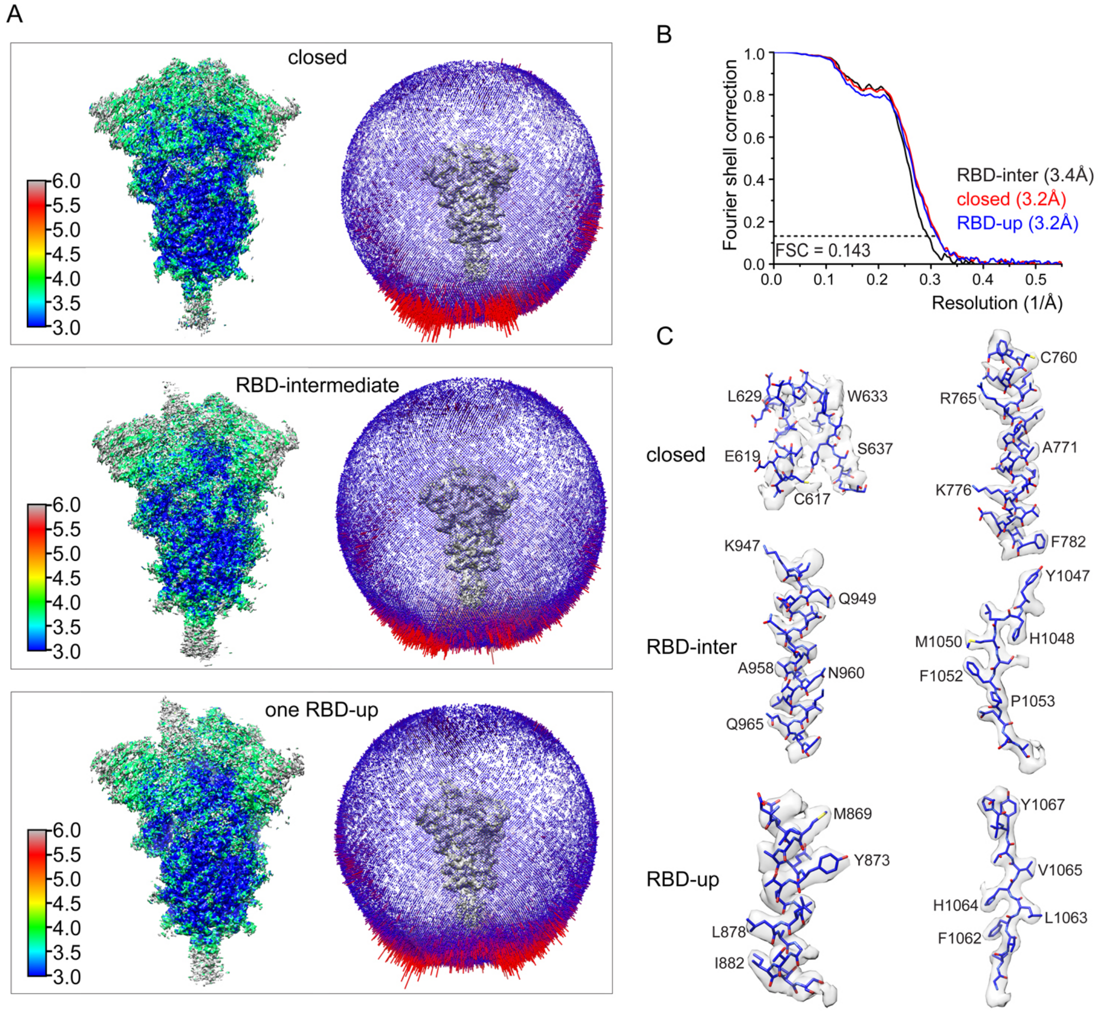


**Figure S3. Analysis of the G614 S trimer structure in DDM.** (A) 3D reconstructions of the G614 S trimer preparation in DDM in the closed, RBD-intermediate and one RBD-up conformations, respectively, are colored according to local resolution estimated by RELION. Angular distribution of the cryo-EM particles used in each reconstruction is shown in the side view of the EM map. (B) Gold standard FSC curves of the three refined 3D reconstructions of the G614 S trimer. (C) Representative density including the 630 loop in gray surface from each EM map.

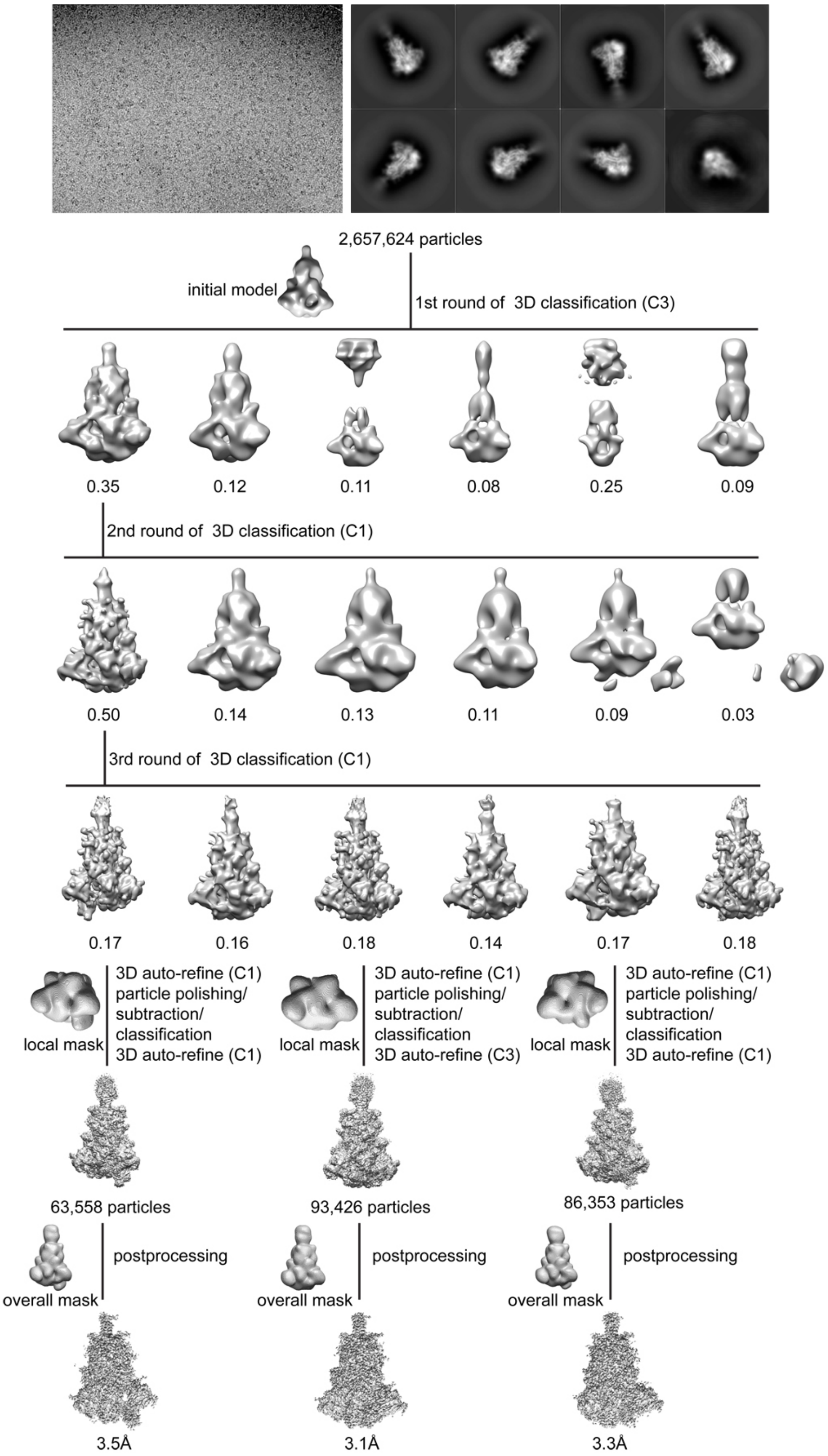
2,657,624 particles
initial model
1st round of 3D classification (C3)
0.35
0.12
0.11
0.08
0.25
0.09
2nd round of 3D classification (C1)
0.50
0.14
0.13
0.11
0.09
0.03
3rd round of 3D classification (C1)
0.17
0.16
0.18
0.14
0.17
0.18
local mask
3D auto-refine (C1)
particle polishing/
subtraction/
classification
3D auto-refine (C1)
local mask
3D auto-refine (C1)
particle polishing/
subtraction/
classification
3D auto-refine (C3)
local mask
3D auto-refine (C1)
particle polishing/
subtraction/
classification
3D auto-refine (C1)
63,558 particles
93,426 particles
86,353 particles
overall mask
postprocessing
overall mask
postprocessing
overall mask
postprocessing
3.5Å
3.1Å
3.3Å

**Figure S4. Cryo-EM analysis of the G614 S trimer in detergent NP-40.** Top, representative micrograph, and 2D averages (box dimension: 396Å) of the cryo-EM particle images of the G614 S trimer in detergent NP-40. Bottom, data processing workflow for structure determination.

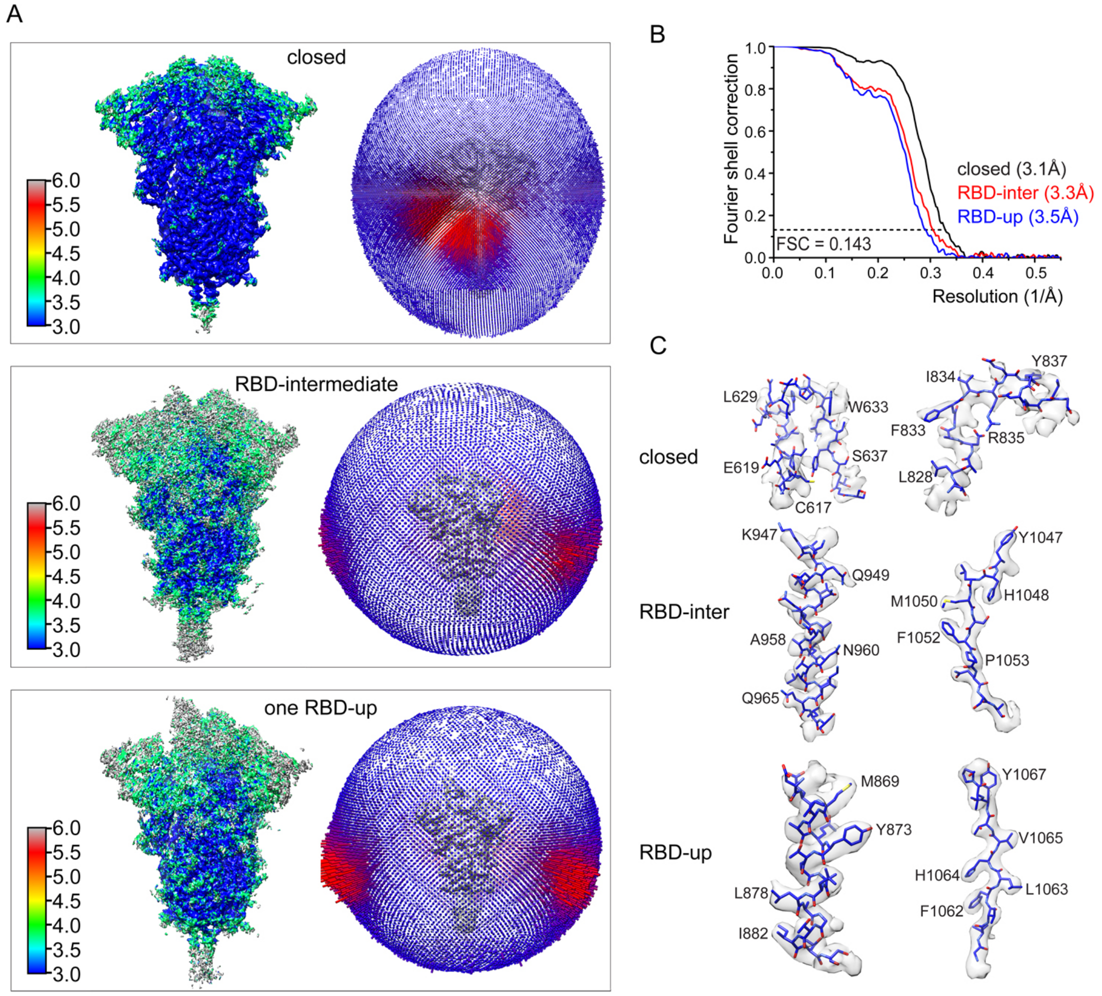


**Figure S5. Analysis of the G614 S trimer structure in NP-40.** (A) 3D reconstructions of the G614 S trimer preparation in NP-40 in the closed, RBD-intermediate and one RBD-up conformations, respectively, are colored according to local resolution estimated by RELION. Angular distribution of the cryo-EM particles used in each reconstruction is shown in the side view of the EM map. (B) Gold standard FSC curves of the three refined 3D reconstructions of the G614 S trimer. (C) Representative density including the 630 loop and FPPR in gray surface from each EM map.

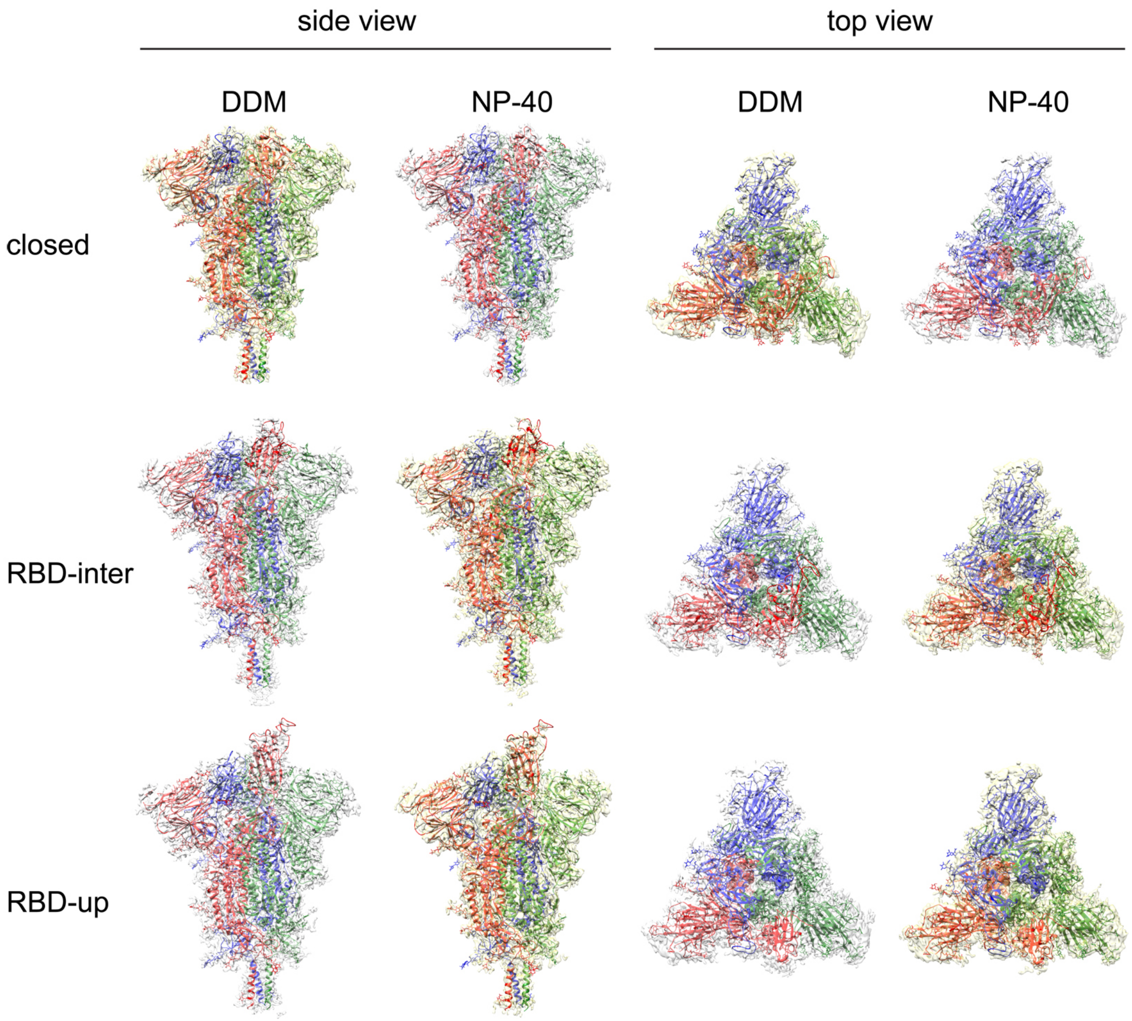


**Figure S6. EM maps for the G614 trimer in detergent DDM and NP-40.** Top and side views of the reconstructions of the G614 trimer in either DDM or NP-40 representing the closed, RBD-intermediate and RBD-up conformations, respectively. The corresponding maps from the two detergents are nearly identical and all maps were used for structural interpretation.

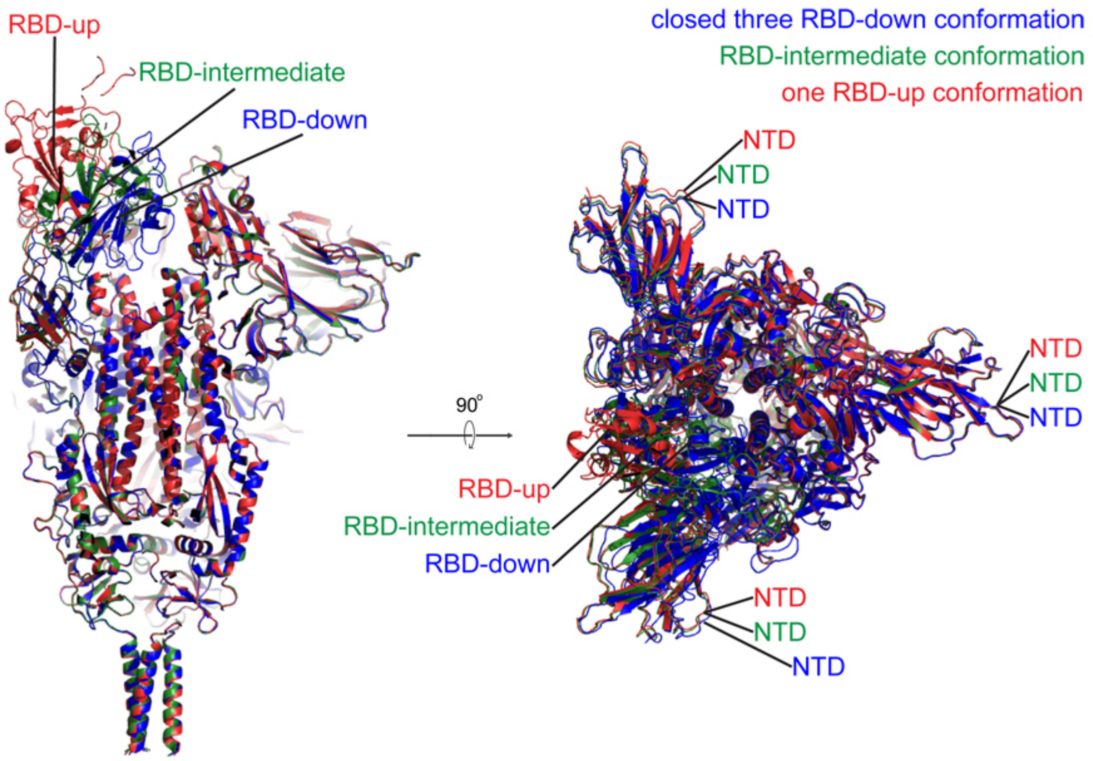


**Figure S7. Superposition of the three structures of the G614 S trimer.** Three structure of the G614 trimer representing the closed (blue), RBD-intermediate (green) and RBD-up conformations (red) are aligned by the invariant S2. Three RBD positions as well as NTDs are indicated.

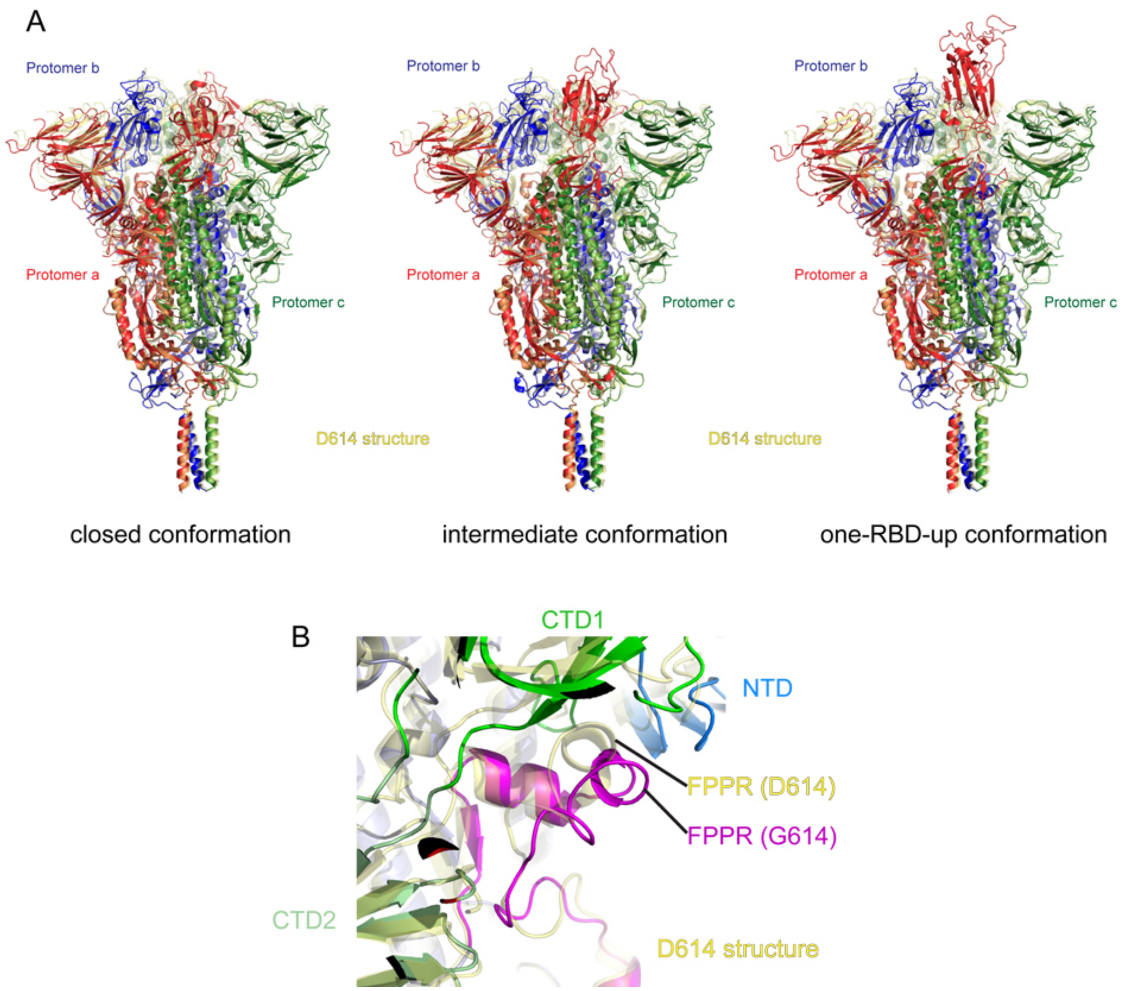


**Figure S8. Superposition of the G614 trimer structures and the D614 structure.** (A) Side views of superposition of three structures of the G614 S in ribbon representation with the structure of the prefusion trimer of the D614 S (PDB ID: 6XR8), shown in yellow. Top views of superposition are shown in Figure 2B. (B) Comparison of the FPPR in the G614 and D614 trimer structures.

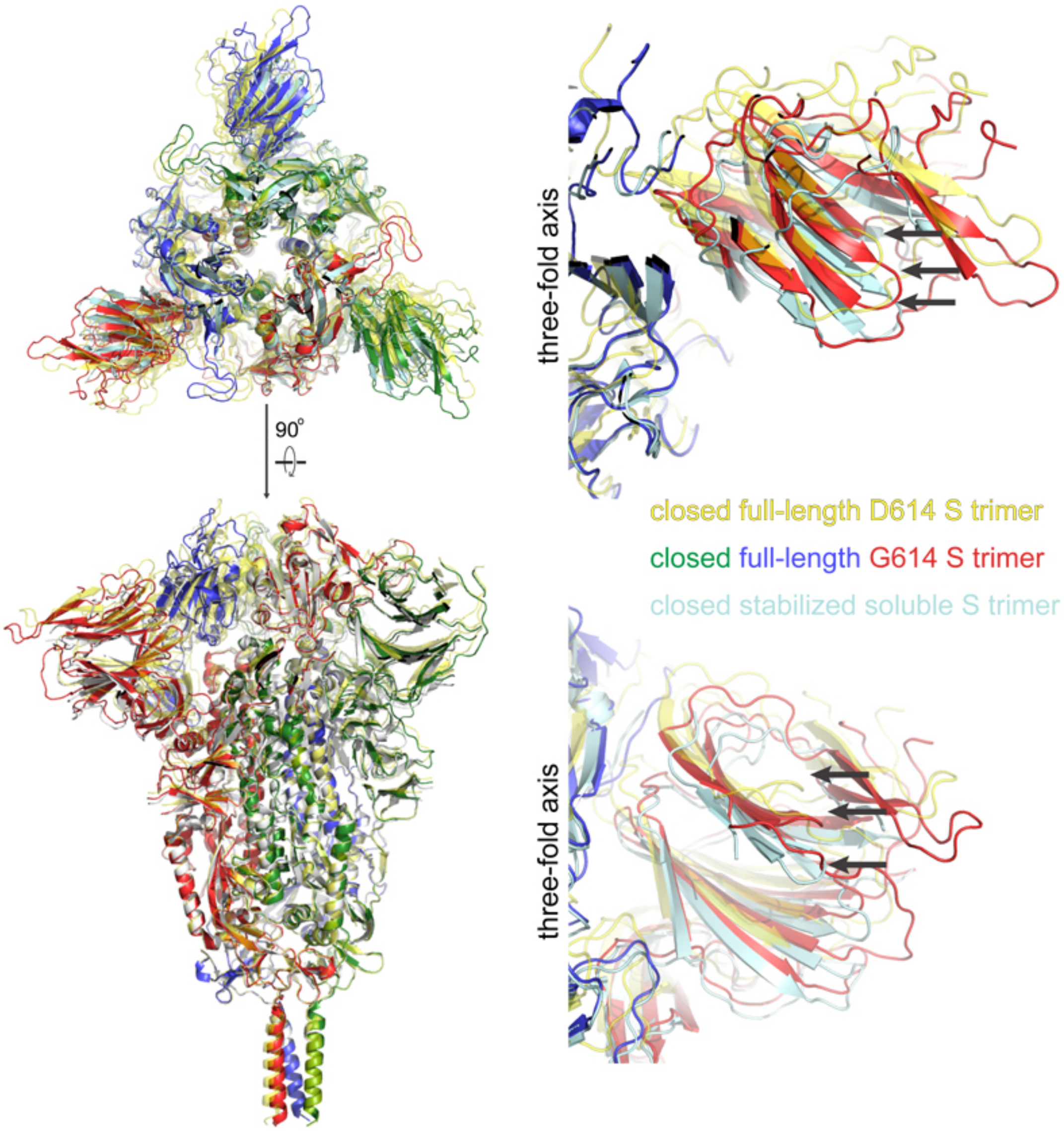


**Figure S9. Superposition of the closed prefusion structures from the full-length G614 trimer, the full-length D614 trimer and the stabilized soluble S trimer.** Three structure of the intact G614 trimer in the closed conformation (red, blue and green), the intact D614 trimer in the closed conformation (yellow; PDB ID: 6XR8 ), and the stabilized S ectodomain in the closed conformation (light blue; PDB ID: 6VXX) are aligned by the invariant S2. Both the top and sides views are shown. Positions of the same loop in the NTD in the three structures are indicated by black arrows.

### Table S1. Binding constants of S-ACE2 interaction

| Construct | KD (M) | KD Error | ka (1/Ms) | ka2 | ka Error | ka2 Error | kdis (1/s) | kdis2 | kdis Error | kdis2 Error |
|---|---|---|---|---|---|---|---|---|---|---|
| D614/monomeric ACE2 | 1.33E-07 | 1.55E-09 | 1.37E+05 | | 1.52E+03 | | 1.81E-02 | | 6.53E-05 | |
| G614/dimeric ACE2 | 3.43E-07 | 8.55E-09 | 7.92E+04 | | 1.90E+03 | | 2.71E-02 | | 1.88E-04 | |
| D614/monomeric ACE2 | 1.12E-08 | 3.28E-10 | 3.95E+04 | 4.80E+00 | 6.85E+02 | 1.40E+01 | 4.42E-04 | 2.77E-01 | 1.05E-05 | 8.00E-01 |
| G614/dimeric ACE2 | 1.55E-08 | 7.11E-10 | 2.98E+04 | 1.31E+00 | 7.50E+02 | 1.59E+00 | 4.62E-04 | 6.79E-02 | 1.77E-05 | 7.92E-02 |

**Table S2. EM data collection and reconstruction statistics**

| Protein | SARS-CoV-2 S prefusion | | | | | |
|---|---|---|---|---|---|---|
| Detergent | NP-40 | | | DDM | | |
| Microscope | Titan Krios | | | Titan Krios | | |
| Voltage(kV) | 300 | | | 300 | | |
| Detector | Gatan K3 | | | Gatan K3 | | |
| Magnification(nominal) | 105,000 | | | 105,000 | | |
| Energy filter slit width (eV) | 20 | | | 20 | | |
| Calibrated pixel size (Å/pix) | 0.825 | | | 0.825 | | |
| Exposure rate ($e^-$/pix/sec) | 14.77 | | | 14.68 | | |
| Frames per exposure | 50 | | | 50 | | |
| Total electron exposure ($e^-$/Å$^2$) | 54.3 | | | 53.9 | | |
| Exposure per frame ($e^-$/Å$^2$) | 1.09 | | | 1.08 | | |
| Defocus range (µm) | 1.5-2.7 | | | 1.0-2.5 | | |
| Automation software | SerialEM | | | SerialEM | | |
| # of Micrographs used | 11,577 | | | 11,092 | | |
| Particles extracted | 3,640,242 | | | 5,652,781 | | |
| Class | closed | inter | open | closed | inter | open |
| Total # of refined particles | 93,426 | 86,353 | 63,558 | 55,096 | 63,040 | 55,537 |
| Symmetry imposed | C3 | C1 | C1 | C1 | C1 | C1 |
| Estimated accuracy of translations/rotations | 1.01/2.0 | 1.28/2.13 | 1.46/2.60 | 1.50/2.58 | 1.24/2.07 | 1.71/2.73 |
| Map sharpening B-factor | -81.6 | -65.5 | -86.3 | -79.1 | -59.2 | -55.2 |
| Unmasked Resolution at 0.5/0.143 FSC (Å) | 3.96/3.47 | 6.6/3.88 | 6.5/3.96 | 6.65/4.2 | 6.49/3.80 | 6.55/3.67 |
| Masked resolution at 0.5/0.143 FSC (Å) | 3.50/3.07 | 3.86/3.33 | 4.0/3.50 | 3.93/3.44 | 3.81/3.21 | 3.82/3.24 |

**Model refinement and validation statistics**

| Class | closed | inter | open |
|---|---|---|---|
| PDB | | | |
| Composition | | | |
| Amino acids | 3363 | 3326 | 3325 |
| Glycans | 57 | 57 | 57 |
| RMSD bonds (Å) | 0.013 | 0.013 | 0.016 |
| RMSD angles (◦) | 1.934 | 1.952 | 2.104 |
| Mean B-factors | | | |
| Amino acids | 47.88 | 47.81 | 47.81 |
| Glycans | 90.03 | 90.18 | 90.07 |
| Ramachandran | | | |
| Favored (%) | 93.74 | 93.42 | 92.96 |
| Allowed(%) | 5.99 | 6.13 | 6.80 |
| Outliers(%) | 0.027 | 0.46 | 0.24 |
| Rotamer outliers (%) | 1.33 | 0.97 | 1.10 |
| Clash score | 8.04 | 6.31 | 6.14 |
| C-beta outliers (%) | 0.29 | 0.19 | 0.71 |
| CC (mask) | 0.83 | 0.79 | 0.79 |
| MolProbity score | 1.95 | 1.78 | 1.82 |
| EMRinger score | 3.69 | 3.40 | 3.26 |
